# Genetic analysis reveals efficient sexual spore dispersal at a fine spatial scale in *Armillaria ostoyae*, the causal agent of root-rot disease in conifers

**DOI:** 10.1101/105825

**Authors:** Cyril Dutech, Frédéric Labbé, Xavier Capdevielle, Brigitte Lung-Escarmant

## Abstract

*Armillaria ostoyae* (sometimes named *A. solidipes*) is a fungal species causing root diseases in numerous coniferous forests of the northern hemisphere. The importance of sexual spores for the establishment of new disease centers remains unclear, particularly in the large maritime pine plantations of southwestern France. An analysis of the genetic diversity of a local fungal population distributed over 500 ha in this French forest showed genetic recombination between genotypes to be frequent, consistent with regular sexual reproduction within the population. The estimated spatial genetic structure displayed a significant pattern of isolation by distance, consistent with the dispersal of sexual spores mostly at the spatial scale studied. Using these genetic data, we inferred an effective density of reproductive individuals of 0.1 to 0.3 individuals/ha, and a second moment of parent-progeny dispersal distance of 130 to 800 m, compatible with the main models of fungal spore dispersal. These results contrast with those obtained for studies of *A. ostoyae* over larger spatial scales, suggesting that inferences about mean spore dispersal may be best performed at fine spatial scales (i.e. a few kilometers) for most fungal species.

## Introduction

During their life-cycle, fungal species may either alternate sexual and asexual reproduction, or reproduce exclusively by one of these two modes of reproduction (e.g. Taylor et al. 1999). These reproduction strategies have important consequences for fungal demographics and evolution. For example, mostly asexual populations can produce large numbers of progenies rapidly, enabling them to colonize new environments to which they are already adapted (e.g. Hovmøller et al. 2008, Raboin et al. 2007). However, the absence of genetic recombination may decrease adaptation potential in a changing environment, because new genotypes can emerge only by mutation (Crow & Kimura 1965). Sexual and asexual propagules may also have different dispersal patterns, generating different population genetic structures at different spatial scales (Barrès et al. 2012, Rieux et al. 2014). These different dispersal capacities may produce a spatial and temporal mosaic of genetic diversity, allowing natural selection to act at different spatial and temporal levels (Peay & Bruns 2014, Thrall & Burdon 2002). Mode of reproduction is, therefore, an important biological trait that may vary with ecological context, as predicted and observed in invasive species (Bazin et al. 2014, Gladieux et al. 2015).

It is not always easy to infer the mode of reproduction and dispersal processes within species, or between populations of the same species. Sexual reproduction may be cryptic (Saleh et al. 2012), or not very efficient at producing new recombining genotypes within populations (Dutech et al. 2010). Direct observations in the field may be rare or inconclusive. Population genetic analyses may therefore be more relevant for estimating the contribution of sexual reproduction events within and between populations (Halkett et al. 2005). First, analyses of genetic diversity performed at different spatial scales may reveal the different reproductive processes within species (e.g. Barrès et al. 2012, Dutech et al. 2008, Kohli et al. 1995). Second, methods using spatial genetic analysis to estimate the genetic relatedness between individuals can be used to infer the spatial range of dispersal of sexual and asexual spores (e.g. Dutech et al. 2008, Rieux et al. 2011, Travadon et al. 2012), under the theoretical model of isolation by distance (IBD) (Rousset 1997, Vekemans & Hardy 2004, Wright 1943). However, despite the common use of such methods on fungal populations, the conclusions drawn about dispersal processes may sometimes be irrelevant, due to the use of an inappropriate spatial scale of investigation (see for example Rousset 1997 for theoretical considerations). Thus, conclusions that long-distance spore dispersal occurs based on an absence of spatial genetic structure among populations may be incorrect for several fungal species, as already shown for marine organisms with a large geographic distribution (Puebla et al. 2012). Puebla *et al*. showed that genetic analyses over fine spatial scales (i.e. a few meters to a few kilometers), and estimates of relatedness between genotypes, could, in some cases, better detect the effect of restricted progeny dispersal than studies performed at population level.

*Armillaria ostoyae* (Romagnesi) Herink is a fungal species for which the importance of sexual reproduction in the life-cycle remains unclear (Rishbeth 1988, Prospero et al. 2008). It was recently proposed an older name for this species: *A. solidipes* Peck, but this replacement name may be incorrect and may create confusion (Hunt et al. 2011). *Armillaria ostoyae*, responsible for butt- and root-rot diseases of conifers throughout the northern hemisphere, has two known modes of reproduction and dispersal. First, *A. ostoyae*, as other *Armillaria* species, produces a specific mycelium, known as a rhizomorph, for dispersal in the soil from one infected root to another susceptible host root (Lung-Escarmant & Guyon 2004, Redfern & Filip 1991). This vegetative propagation, also associated with a direct transmission from root to root, may lead to the development of a single-genotype clonal patch in a forest stand, sometimes covering a very large area (up to about 1000 ha; Ferguson et al. 2003). *A. ostoyae* also regularly produces fruiting bodies during the fall, and these fruiting bodies can release sexual haploid spores (i.e. the basidiospores) into the air. The germination of two haploid compatible basidiospores and the fusion of their hyphae give rise to a new diploid genotype able to infect a new host tree (Rishbeth 1988). It has been assumed that sexual spores may occasionally be dispersed, by the wind, over tens of kilometers in many fungal species (Aylor 1990, Halbwachs & Bässler 2015). This mode of reproduction should, therefore, facilitate colonization by new genotypes and the development of new clonal patches some distance away from the original disease focus. However, the fusion of two sexual basidiospores leading to the colonization of a new host has rarely been observed or achieved in the field for *A. ostoyae*, raising questions about the frequency with which this process occurs in natural conditions for this species (e.g. Rishbeth 1970, Rishbeth 1988).

Several studies on *A. ostoyae* across the northern hemisphere have identified different population genetic structures potentially dependent on the ecological and climatic context of the forests studied. For example, in extensively managed cold North American forests, clonal structures of several tens to hundreds of hectares in size have frequently been reported (e.g. Dettman & Van der Kamp 2001a, Fergusson et al. 2003). By contrast, in the conifer forests of European mountain ranges, in which human activities have had a stronger impact on the ecosystem over a longer period of time, *A. ostoyae* generally displays higher levels of genotypic diversity and forms smaller clonal patches (e.g. Bendel et al. 2006, Legrand et al. 1996). In the maritime pine plantations of southwestern France, the Landes de Gascogne forest, small clonal patches associated with disease foci have also been described (Prospero et al. 2008). This genetic structure would be expected for a forest growing in a warm and wet climate, and with a rapid turnover of susceptible hosts due to intensive management of the plantation (Fergusson et al. 2003, Rishbeth 1988). As in previous studies, no genetic relatedness was detected between clonal patches. Two mutually exclusive hypotheses can be put forward to explain this absence of genetic structure. The first hypothesis is that sexual spores are regularly dispersed over long distances (i.e. over several tens of kilometers), with efficient germination and fusion of basidiospores, in this forest, resulting in a random genetic structure (Prospero et al. 2008). This hypothesis has been also proposed for *A. mellea* in North America to explain the low estimated population genetic differentiation (Baumagartner et al. 2010). However, this hypothesis is not consistent with the limited spread of the disease observed over several decades in this forest. The disease is thought to have originated on the west coast of the region, where most forest areas were located before the large maritime pine plantations were established in the 19^th^ century (Labbé et al. 2015, Prospero et al. 2008). The second hypothesis is that the spatial scale of the previous study, covering the entire area of the forest (i.e. around 1 × 10^6^ ha) was inappropriate. Rare long-distance dispersal events can occur for sexual spores, but the mean dispersal distance is probably in the order of a few kilometers, as observed in other fungi (Rieux et al. 2014). In this context, as highlighted above, genetic studies at a spatial scale similar to that over which dispersal occurs are required to detect any spatial genetic structure associated with the dispersal process. We tested the IBD hypothesis at local scale, by analyzing the spatial genetic structure of *A. ostoyae* within a coastal forest in which several genotypes were identified in a previous study (Prospero et al. 2008), expanding the sampled area from less than one hectare to several hundred hectares.

Our main objectives were to use the recently developed molecular markers for *A. ostoyae* (Dutech et al. 2016) to: 1) estimate the importance of genetic recombination in the population studied; 2) characterize the spatial genetic structure of this population; 3) test the hypothesis of isolation by distance (Wright 1943), assuming that the mean dispersal distance of sexual spores is between a few hundred meters and a few kilometers; 4) estimate the spatial range over which spores are dispersed from genetic data.

## Materials and Methods

### Study site

The study was performed in the western part of the Landes de Gascogne Forest, in La Forêt Domaniale de Saint Julien en Born (1°,16′,20″W – 44°,06′,40″N) in southwestern France. This site is located on sandy dunes 4 km from the Atlantic Ocean. The forest is composed mostly of maritime pines (*Pinus pinaster*), with some oaks (*Quercus ilex, Q. robur)*. Maritime pine is a local species that has been cultivated intensively in plantations since the 19^th^ century, to help drain the marshes and swamps initially present in the area and for economic reasons (for details, see Labbé et al. 2015). The populations of maritime pine on the coast were present before these large plantations were established. They formed small fragments of forest and were the first populations sown during this period to stabilize the sandy coastal dunes (Labbé et al. 2015). The coastal part of the Landes de Gascogne Forest is currently managed by the French National Forestry Office (*Office National des Forêts*), which favors management by natural regeneration or the sowing of maritime pine seeds after commercial logging.

We extended the initial sampling area used by Prospero et al. (2008), from the disease focus named “Contis” to the south of the stand. We obtained 177 mycelial fans from dead or dying maritime pines, by collecting mycelium from under the cork at the collar of the trees as described by Prospero et al. (2008). This previous study clearly identified clonal patches that were generally less than 1 ha in diameter and associated with a single disease focus. We maximized the chances of collecting different genotypes for studies of their genetic relationships, by collecting samples mostly from trees located at least 100 m apart (55% of the samples). The minimum distance separating two samples was 16 m; the maximum distance was 3474 m, and the total area sampled was close to 500 ha. Samples were located with a GPS Trimble Geo7X, and GPS Pathfinder Office (Trimble Navigation Ltd, USA) was used for the processing of spatial data.

### Molecular analysis

In the laboratory, we collected mycelium fans from each infected piece of wood sampled in the field, by detaching the mycelium from the cambium with a scalpel. These mycelium were freeze-dried overnight (-45°C, 0.3 mbar), and then ground with metallic beads in 2 ml microtubes, with an automatic grinder (GenoGrinder Sample Prep 2010, SPEX, USA), for 15 s at 1500 rpm. DNA was extracted from the ground mycelium in a cetyltrimethyl ammonium bromide (CTAB) buffer, according to the protocol described by Prospero et al. (2008).

Samples were genotyped for 27-single nucleotide polymorphisms (SNPs) identified in 24 genes present as single copies in most fungal genomes, and isolated by the PHYLORPH method (see for details, Feau et al. 2011). These 27 SNPs were selected from the list of 82 SNPs previously validated in two coastal *A. ostoyae* populations (Dutech et al. 2016). These SNPs were multiplexed, and genotyped in the medium-throughput MassARRAY iPLEX genotyping assay from Sequenom (San Diego, CA, USA), as described by Chancerel *et al*. (2013) and Dutech et al. (2016). The choice of these SNPs was the best combination between keeping minimum one SNP per gene and the compatibility of their primers for genotyping them in a single Sequenom run. Details of the 27 genotyped SNPs are provided in Table 1.

**TABLE 1:**
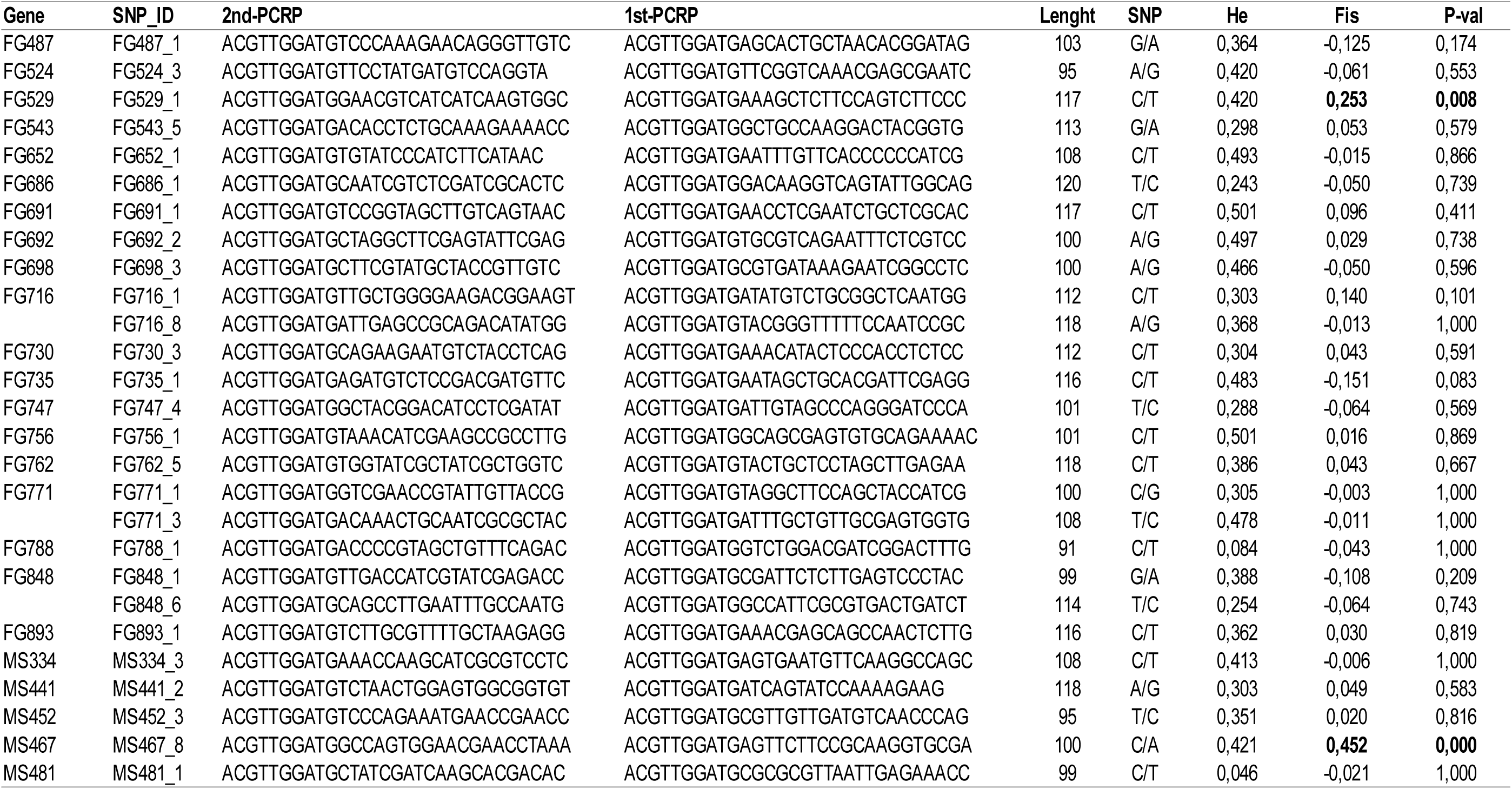
Details of the 27 single nucleotide polymorphisms (SNPs) analyzed in the *A. ostoyae* population sampled in the south-western French forest of St Julien en Born. Gene and SNP_ID (the ID of the analyzed SNP for each gene) refer to the gene and SNP identification given in Dutech et al. (2016). 1rst- and 2ndPCRP refer to the primers used for the Sequenom PCR reaction for each SNP. Length and SNP are respectively the size of the PCR product and the type of nucleotide variation. He and Fis are the gene diversity (Nei, 1987) and the intra-individual genetic fixation index respectively. P-val are the probabilities of the exact test for Hardy-Weinberg expectations (Raymond & Rousset 1995). Values significantly different from a panmictic population are in bold.

### Data analyses

Repeated genotypes (i.e. identical for all 27 SNPs analyzed) were identified with GenoType V.1.2, excluding missing loci for each comparison (Meirmans & Van Tienderen 2004). For each SNP, allelic frequencies, genic diversity (*H_E_*, Nei 1987) and intra-individual fixation index (*F_IS_*) according to the Weir and Cockerham (1984) method were estimated with Genepop V.4.3 (Raymond & Rousset 1995). Only one copy per genotype was retained, to exclude the effect of clonal structure on the estimates of genetic diversity. Exact tests for Hardy-Weinberg equilibrium within each population, and for linkage disequilibrium between SNPs were performed with Genepop. Correction for multiple testing was performed according to the false discovery rate method (FDR, Benjamini & Hochberg 1995), with the R package fdrtool V1.2.15 (Klaus & Strimmer 2013).

Spatial genetic structure was analyzed by two methods. First, the kinship coefficient was estimated for each pair of mycelial samples, using the estimate proposed by Loiselle et al. (1995). The mean per spatial distance classes was then plotted to generate a spatial autocorrelogram of the genetic relationships between samples. We defined the distance classes so as to obtain an even number of pairs of samples in each distance class. The first five distance intervals were each 150 m wide. Beyond 750 m, each of the next five classes was 250 m wide and then, after 2000 m, there were 500 m-wide classes, making it possible to estimate kinship coefficients over distances of up to 3000 m. Each distance class contained at least 210 pairs of samples (obtained for the first distance class with the genotypes in single copy). Random permutations of the spatial locations of samples (1000 permutations) were performed to test the hypothesis that the mean kinship coefficient for each distance class was significantly different from that expected for a random spatial genetic structure. We investigated the effect of clonality on spatial genetic structure, by performing the analysis both with all genotypes and with a single copy of each genotype randomly chosen from the copies.

The second method used to investigate spatial genetic structure was spatial principal component analysis (sPCA, Jombart et al. 2008). This method can be used to identify various spatial genetic structures (e.g. patches, clines, genetic barriers), without the need for assumptions concerning the genetic model, whereas Bayesian genetic clustering methods are dependent on the assumption of Hardy-Weinberg equilibrium. sPCA can be used to investigate the distribution of allelic diversity, by combining a principal component analysis (PCA) of allele frequencies and the spatial distance between samples. It can identify both global structures associated with the positive components of the analysis, reflecting the decrease in similarity between individuals with increasing spatial distance, and local structures associated with the negative components reflecting local dissimilarities between individuals located close together spatially (Jombart et al. 2008). Each significant structure detected was displayed by plotting the samples according to their geographic coordinates, color-coding and sizing them according to their scores along the significant sPCA axes. We determined whether the global and local structures were significantly different from a random structure by performing random permutations of sPCA components (999 permutations), and comparing the observed and simulated sPCA statistics as described by Jombart et al. (2008). Estimation and testing were performed with the R package Adegenet V2.0.1 (Jombart 2008), using a single copy per genotype to eliminate the effect of clonal structure on the analysis.

We tested the hypothesis of isolation by distance (IBD) associated with limited dispersal of the progeny from the parents (Wright 1943), by assuming a spatial dispersal of sexual spores in two dimensions. Under this hypothesis, the slope of the estimated kinship coefficient for each pair of samples decreases linearly with the logarithm of the spatial distance separating these pairs (Rousset 2000). The significance of the slope was tested by permutations of spatial distances between samples (1000 permutations) and Spearman’s rank correlation analysis, as described by Rousset (2000). All the estimations of kinship coefficient and tests were performed with SPAGeDi V.1.5a (Hardy & Vekemans 2002).

We also estimated the effective size (*Ne*) of the *A. ostoyae* population by a method based on the linkage disequilibrium between loci (Do et al. 2014, Waples & Do 2010). Estimates of *Ne* may be biased by a Wahlund effect in continuous populations, due to the limited dispersal of individuals or their gametes (Neel et al. 2013). We therefore estimated this parameter for both the whole sample, with a single copy per genotype, and for three subsamples of 30 genotypes each, with a maximum distance of 1000 m between samples from the same subsample. These subsamples, and which were smaller than the total sample, may be closer to the window in which most mating events occur, for which the effective number of parents responsible for producing the sample is best estimated (Neel et al. 2013). These three subsamples were evenly distributed over the sampling site, from north to south. *Ne* was estimated with NeEstimator V.2.1 (Do et al. 2014).

Finally, we estimated the second moment of parent-progeny dispersal (Rousset 2000) with the iterative procedure implemented in SPAGeDi V.1.5a (Hardy & Vekemans 2002). Assuming gene dispersal in two dimensions and a genetic drift-dispersal equilibrium (Wright 1943, Rousset 2000), the slope of the spatial autocorrelation of the kinship coefficient is 1/(4πD_e_σ^2^), where D_e_ is the effective density of reproductive individuals, and σ is the second moment of parent-progeny dispersal. However, this relationship should only be estimated in the range from σ to around 10 or 20σ (Rousset 2000). Outside this range, it may be biased by the shape of the dispersal function, mutation and migration events. As this range is generally not known, the iterative procedure uses an estimate of D_e_, and yields a first estimate of σ obtained from all the spatial distances analyzed. The value of σ is then re-estimated, considering a spatial range associated with this first estimate of 10σ or 20σ, until σ and the spatial range converge. For D_e_, we used the density of genotypes obtained from our sampling, or the estimate obtained from NeEstimator V.2.1.

## Results

In total, 177 samples were analyzed for the 27 SNPs, and 149 different genotypes were detected. Seventeen genotypes occurred at least twice (eight genotypes occurred twice, seven occurred three times and two occurred four times). The minimum and maximum distance between identical genotypes were 27 m and 371 m, respectively, with a mean value of 120 m. Genetic diversity, estimated with *H_E_*, was between 0.046 (MS481_1) and 0.501 (FG691_1 and FG756_1), with a mean value of 0.36 (SE ± 0.02) (Table 1). The intra-individual fixation index (*F_IS_*) was estimated at between -0.151 (FG735_1) and 0.452 (MS467_8), with a mean value of 0.02 (SE ± 0.02). Overall, genotype frequencies did not differ significantly from that expected under Hardy-Weinberg equilibrium. Two loci (MS467_8 and FG529_1) displayed a significant departure of genotype frequencies from Hardy-Weinberg expectations (*P*-values < 1 × 10^−4^ and 0.008, respectively; Table 1). However, this departure from expectations remained significant only for MS467_8 after correction for multiple tests. In total, 351 comparisons were made for linkage disequilibrium, and 37 pairs of loci had significant *P*-values (< 0.05). After correction for multiple testing, only four tests remained significant. Two were associated with genes from the same locus (FG771_1 and FG771_3, and FG848_1 and FG848_6). The other two significant tests concerned FG730_3 and MS452_3, and FG893_1 and MS334_3. Finally, to avoid any effect of linkage disequilibrium or departure from Hardy-Weinberg equilibrium on demographic inferences, we removed MS467_8, FG771_1, FG848_1, FG730_3 and FG893_1, and used only 22 SNP loci for subsequent analyses.

The estimated kinship coefficient between pairs of *A. ostoyae* samples decreased steadily with increasing distance (Figure 1). Taking all the samples into account, the estimated kinship coefficient was 0.09 for the first distance class (0-150 m) and was close to zero beyond 450 m. With the exception of two distance classes (1500-1750 m and 1750-2000 m) kinship coefficient estimates at distances of more than 450 m were not significantly different from expectations for a random genetic structure. For the two remaining distance classes, the kinship coefficient was estimated at -0.01 and -0.025, respectively. A similar decrease in kinship coefficient with increasing distance was observed when only one copy of each genotype was considered. The estimate was lower for the first distance class (0.04), but identical for the second class and beyond, and not significantly different from that expected under a random genetic structure, except, once again, for the 1500-1750 m and 1750-2000 m distance classes. The slope of the autocorrelation for one copy per genotype was – 0.012 (SE ± 0.003), which was significantly different from zero in Spearman’s rank correlation analysis on spatial locations (*P-value* < 0.01).

**Figure 1:**
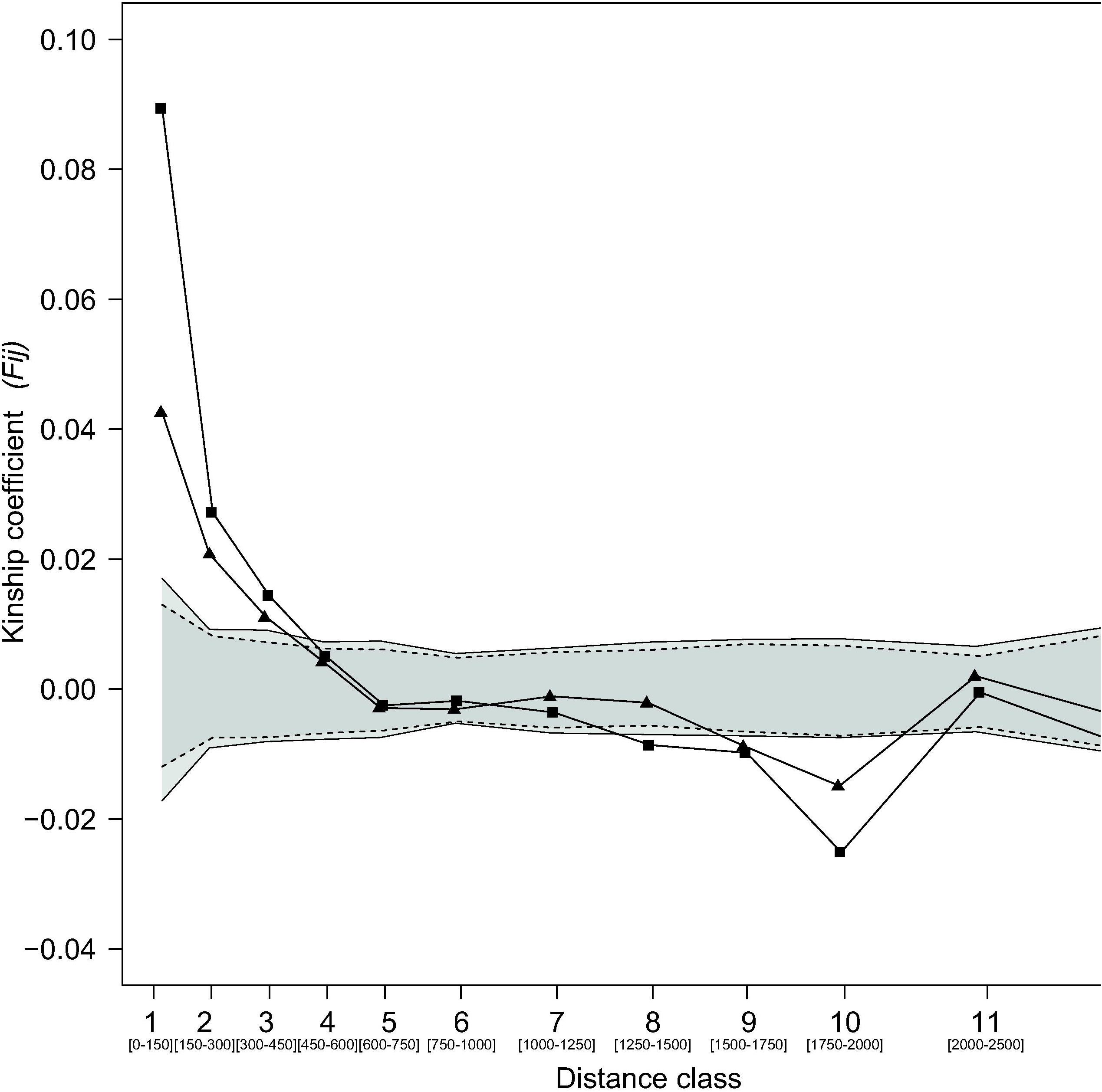
Relationships between the kinship coefficients of pairs of *A. ostoyae* samples (*F*_ij_) and the distance class separating them. Range of each class was indicated in meters in bracket. Square symbols represent the kinship coefficients estimated using all the samples, and triangular symbols represent those estimated from only one copy per genotype (i.e. clonal correction). The light gray area corresponds to 95% of the expected kinship coefficient assuming a random spatial genetic structure for all samples considered, and the dark gray area corresponds to that based on only one copy per genotype.

Tests on the spatial structure estimated from sPCA showed that only the global structure, associated with positive eigenvalues, was significantly different from a random spatial structure (999 permutations, *P*-value = 0.011). The local structure associated with negative eigenvalues (999 permutations, *P*-value =0.256) was not significant. Several spatial clusters of samples with either the most positive or the most negative values on the first principal axis were observed within the population (Figure 2). Samples with a value close to zero were generally located between these clusters. These spatial clusters had a diameter of about 500 m-1000 m (Figure 2).

**Figure 2:**
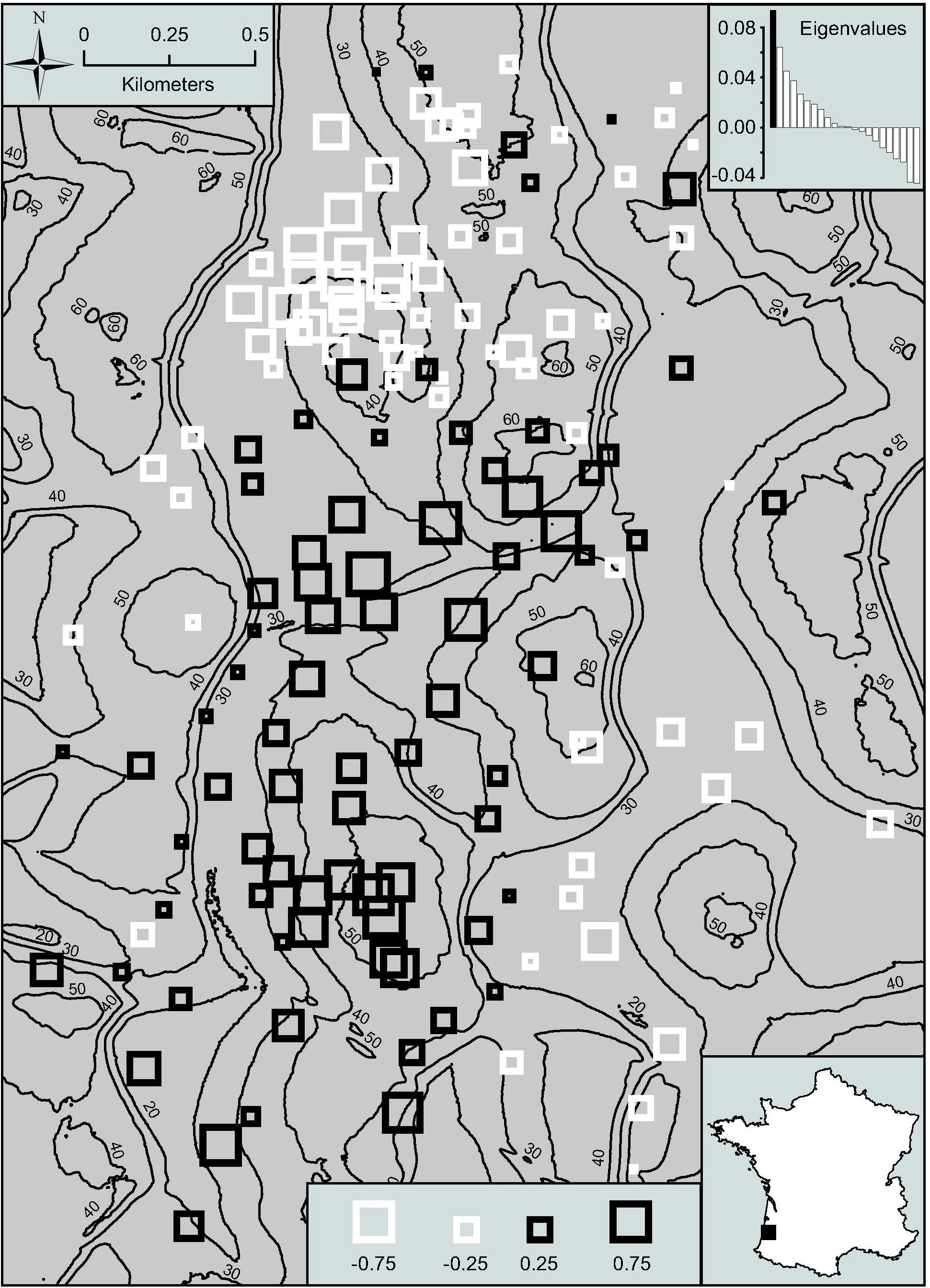
Geographic distribution of individual scores on the first positive axis of the sPCA for the *A. ostoyae* population of southwestern France. Positive and negative values are shown in black and white, respectively, and the size of the squares is proportional to their absolute value. The eigenvalues of each sPCA axis are presented at the top right of the figure, and the geographical location of the site study in France at the bottom right.

For the 149 genotypes sampled from an area of about 500 ha, the *Ne* estimate was 49.9 (95% CI, determined by a jacknife procedure: 33.6 – 77.3), yielding a D_e_ of 0.1 parents/ha. For each subsample of 30 genotypes from about 100 ha, the estimated value of *Ne* ranged between 14.4 and 79.6 (mean = 40; D_e_ = 0.4 parents/ha). We therefore used three effective densities of parents to estimate σ, with the iterative procedure implemented in Spagedi. The first density used was that estimated with NeEstimator V.2.1 and the 149 genotypes. The second was the density of sampled genotypes over an area of 500 ha (0.3 parents/ha, similar to the density obtained with the three subsamples and NeEstimator V.2.1), and the third was a density 10 times greater than the census density, assuming an incomplete sampling of the area (3 parents/ha). The estimated value of σ was between 130 and 162 m for the highest effective density of 3 parents/ha, depending on the maximum distances considered for the regression (10 σ and 20 σ, respectively). The corresponding estimate was 830 m for the sampled density. No convergence was observed for density estimates obtained for the 149 genotypes with NeEstimator.

## Discussion

The observed clonal fraction in this study was lower than previously reported for other European and North American *A. ostoyae* populations (e.g. Bendel et al. 2006, Dettman & Van Der Kamp 2001a, Prospero et al. 2003). In this forest in southwestern France, 90% of the genotypes identified were sampled only once. By contrast, in most of the other forests studied to date, most, if not all, genotypes were sampled several times. The sampling design of this study, with a minimal distance of 100 m between most of the pairs of field samples, may account for the small number of repeated genotypes. In a previous study on this maritime pine forest, most of the identical genotypes associated with a single disease focus were sampled from trees less than this minimal distance apart (Prospero et al. 2008). It was difficult to estimate the size of clonal patches from our sample, but our results are consistent with an area of less than 1 ha, the estimate obtained in a previous study on this forest (Prospero et al. 2008, Labbé 2015, Lung et al. unpublished). This mean area of disease foci is among the smallest clonal patch sizes reported for *A. ostoyae* populations in Europe and North America, where estimates of greater than 1 ha, and up to 965 ha have been obtained (Bendel et al. 2006, Dettman and Van Der Kamp 2001a, Ferguson et al. 2003).

The large number of different genotypes estimated to be present in an area of only 500 ha in this study may be due to the more frequent production of fruiting bodies in coniferous forests in temperate climates than in those growing in a boreal of alpine climate. Wet climates, as for this European coastal area, are assumed to be more conducive to *Armillaria* reproduction and germination of basidiospores on stumps or pieces of wood (Rishbeth 1970). This higher frequency of sexual reproduction would favor the establishment of new genotypes over time, through the fusion of sexually compatible basidiospores, resulting in the establishment of large numbers of small clonal patches at the local scale. This hypothesis has been put forward and discussed before (Ferguson et al. 2003, Wargo & Shaw 1985), but there is currently no clear evidence for such a variation of fruiting in *A. ostoyae* populations with temperature and humidity. A second factor potentially explaining the higher level of local genotypic diversity is the extinction-recolonization dynamics of genotypes associated with high levels of environmental disturbance. In intensively managed forests, such as this single-species plantation, successive planting events after the removal or destruction of stumps could eradicate clonal patches, or at least initially decrease their size (Cleary et al. 2013). However, subsequently, as suggested in several previous studies, clear-cutting and the planting of new stands of conifers would favor colonization by new genotypes from basidiospores (see Rishbeth 1988, Worrall 1994, Legrand et al. 1996). Fresh dead wood remaining after logging, such as stumps, or the small pieces of root remaining in the soil after stump removal, could serve as a substrate for the germination and fusion of basidiospores from neighboring infected forest stands (Rishbeth 1970, Rishbeth 1988), although this remains to be demonstrated. Plantations of maritime pine seedlings, which are very sensitive to *A. ostoyae* in the first years of their life (Lung-Escarmant & Guyon, 2004, Labbé et al. 2015), would also be favourable environments for the rapid spread of these new genotypes. Furthermore, the coastal region in which this study was performed, would have supported a large fungal population for several hundred years, potentially accounting for the large number of genotypes observed. Maritime pine populations have been present in this area for centuries and they may have hosted a large *A. ostoyae* population, consistent with the substantial presence of disease due to this fungus observed in this coastal area (Labbé et al. 2015). In addition, the first seeds of this tree species were sown at this site, to stabilize the sandy dunes, in the middle of the 19^th^ century, and this site has been managed ever since so as to keep the coastal forest intact, thereby decreasing the risk of local extinction for fungal pathogen populations.

A role for basidiospores in the establishment of genotypes was strongly suggested, but not clearly demonstrated, in a previous study on *A. ostoyae* in this planted forest of maritime pine (Prospero et al. 2008). The results reported here provide even stronger support for the hypothesis that the genotypes observed originated from the local dispersal of basidiospores rather than the evolution of divergent clonal lineages present for long periods in the soil, as is sometimes assumed (e.g. Worall et al. 1994). We observed no strong genetic linkage disequilibrium among the SNP loci, consistent with populations evolving sexually, at least occasionally, in the last few years (Halkett et al. 2005, Tibayrenc et al. 1991). Furthermore, a long history of clonal evolution should produce a Meselson effect, with allelic divergence leading to high levels of heterozygosity within genotypes (e.g. Ali et al. 2014). Such an effect is inconsistent with the observed level of homozygosity in this study, which was not significantly different from that expected. In addition, the steady decrease in genetic relatedness with increasing spatial distance between the pairs of samples observed on the spatial correlogram is consistent with the IBD model, and the local dispersal of basidiospores, producing new genotypes. We cannot rule out the possibility that basidiospores can also fuse with some genotypes already present in the stand, via Buller’s phenomenon, to produce new genotypes (Rizzo & May 1994). However, even if this process has been described in the laboratory, there is no evidence to suggest that it occurs frequently in natural conditions.

The presence of these recombining genotypes in this French population suggests that sexual reproduction is a key process in the population dynamics of *Armillaria* species. A similar genetic IBD pattern was detected for an *A. mellea* population in California (Travadon et al. 2012), and the authors also concluded that the local dispersal of basidiospores played an important role in the observed spatial genetic structure. By contrast, although numerous genotypes were observed, no IBD pattern was identified for *A. cepistipes* at nationwide scale in Switzerland (Heinzelmann et al. 2011), and a similar lack of IBD was also observed by Prospero et al. (2008) for *A. ostoyae* at the scale of this French forest region. These conflicting results for IBD pattern highlight the importance of choosing an appropriate spatial scale for inferences concerning the spatial dispersal of progenies based on population genetic methods. If the spatial scale is too large relative to mean dispersal distance, the spatial genetic structure may be missed due to the complex patterns associated with the random effects of rare long-distance dispersal events, and genetic methods may not detect these processes efficiently (e.g. Schwartz & McKelvey. 2009, Wingen et al. 2007). If the spatial scale is too small relative to the mean dispersal distance, there may be too few comparisons between genotypes, which may constitute a serious limitation to the detection of spatial genetic structures, especially after removal of the clonal structure (e.g. Dutech et al. 2008). For example, the use of too few comparisons may account for the random structure observed in a North American population of *A. ostoyae*, in which only a few transects of a few hundred meters in length were studied (Dettman & Van der Kamp 2001b).

The IBD pattern observed in this study was evenly distributed over the site, as sPCA revealed several spatial clusters of related genotypes with a diameter of about 1 km. This patchy genetic structure is consistent with an IBD pattern (Jombart et al. 2008), and a dispersal of the sexual spores over the spatial range of a few kilometers. This result suggests that there was no significant genetic structure, such as a genetic cline or barrier, other than IBD at this spatial scale. Spatial Bayesian clustering analyses detecting no significant genetic cluster also yielded similar conclusions (results not shown). Assuming that the *A. ostoyae* population is close to the genetic drift-dispersal equilibrium, then the genetic structure observed in this study can be attributed principally to IBD, and gene flow can be estimated from the slope of the spatial autocorrelation (Rousset et al. 2000, Vekemans & Hardy 2004). In the coastal area, the assumption of genetic equilibrium for *A. ostoyae* populations is realistic, because maritime pines were present for many thousands of years before the establishment of the large pine plantations in the 19^th^ century (Paquereau 1964). Therefore, with an estimated generation time of between 10 and 20 years (Labbé 2015), several hundreds of generations of the fungus have probably developed in this area. Furthermore, simulations have shown that the IBD pattern rapidly becomes established at fine spatial scales after tens of generations for small effective population sizes, such as that estimated here (Bradbury & Bentzen 2007, Leblois et al. 2003).

Assuming dispersal-drift equilibrium, we tried to estimate the second moment of parent-offspring dispersal, using several estimates of the effective density of parents. The method used to estimate this density yields robust estimates in isolated and panmictic populations, but has some biases when applied to continuous populations with an IBD structure (Neel et al. 2013). For example, when the sampling window is larger than the breeding window (i.e. “the local area where most matings occur”, Neel *et al*. 2013), the population size may be underestimated for the sample. This estimated population size is often only one tenth the effective population size of the sampled area (Neel et al. 2013). Finally, we obtained estimated densities close to 0.3 parents/ha for the two types of sampling (i.e. total sample and subsamples), and close to the number of sampled genotypes. These similar results suggest that this estimate for this *A. ostoyae* population is relatively robust, and that the contribution of sexual reproduction is similar for each of the 149 genotypes sampled in this area. Only a small number of studies have reported estimates of effective population size for fungal populations. Ali et al. (2014) obtained an estimate of between 30 and 40 individuals for several Pakistani *Puccinia* populations, based on temporal genetic sampling. These results, which are similar to our own, indicate that effective population size may not always be as important as frequently assumed (e.g. McDonald & Linde 2002). Using this estimate of effective population size and the iterative procedure, we obtained a second moment of parent-descendant dispersal distance of between 100 and 800 m. This spatial range of dispersal is consistent with direct estimates of the mean dispersal distances for spores obtained for other fungal species and by direct methods (e.g. Rieux et al. 2014). All these results strongly suggest that long-dispersal events do occur occasionally, as reported for many other fungal species (Barrès *et al*. 2008), but that spore dispersal is mostly local (over a few hundred meters). This estimate of limited dispersal suggests that ecological and evolutionary dynamics of *Armillaria* populations mainly occur at landscape scale, as for many microbial species (Peay and Bruns, 2014).

It is generally not possible to draw any firm conclusions about the effect of basidiospore dispersal on the genetic structure of *A. ostoyae* populations from direct observations (e.g. Rishbeth 1988). By contrast, our study clearly shows that, for this species, as for many other fungi, an analysis of spatial genetic structure can provide estimates of both sexual reproduction within populations and the mean distance over which basidiospores disperse. However, we found no evidence of the significant spatial genetic structure observed for this species at larger spatial scales in this forest area (Prospero et al. 2008, Labbé 2015, Dutech et al. 2016). Our results are similar to those obtained for maritime organisms, for which non-linear patterns of IBD have been observed, depending on the spatial scale studied (Bradbury & Bentzen 2007). These studies have revealed that, for organisms with large spatial distributions, such as many fungal species, limited dispersal between the progenies and their parents is often estimated at fine spatial scales, but not at large spatial scales. Rather than being associated with high levels of gene flow that are difficult to estimate with genetic methods at large spatial scales (Wingen et al. 2007), these contrasting results reveal that choosing the most appropriate spatial scale of investigation is crucial to the correct estimation of dispersal processes by genetic methods, as pointed out by Rousset (1997). Conclusions about long-distance spore dispersal may, therefore, often be incorrect for many fungal species if only genetic differentiation between populations is considered. We advocated estimating dispersal at different spatial scales, to avoid misleading conclusions likely to hinder our understanding of fungal biology.

## Acknowledgments

We would like to thank M. Martin, N. Leymarie and O. Fabreguettes for their support and assistance during DNA extraction and laboratory work, and J-P Soularue for his useful comments. Genotyping was performed at the Genomic and Sequencing Facility of Bordeaux (grants from the *Conseil Regional d’Aquitaine* no. 20030304002FA and 20040305003FA and from the European Union, FEDER no. 2003227 and from *Investissements d’avenir, Convention attributive d’aide* No. ANR-10-EQPX-16-01). This work was supported by the POURPIN project (*Ministère français de l’agriculture et de la forêt, Département Santé des Forêts)* and by European funding through the interregional SUDOE FORRISK project (Network for Innovation in Silviculture and Integrated Risk Management Systems in the Forest). F. Labbé was supported by a grant from INRA/Région Aquitaine.

